# Ume6 Coordinates Lysosomal Remodeling and Autophagy Through Transcriptional Repression in *S. cerevisiae*

**DOI:** 10.64898/2026.10.29.762757

**Authors:** Olivia Evans, Alexis Hoy, Isabel Woo, Xi Yang

## Abstract

Lysosome function and autophagy are highly regulated processes that must be coordinated to enable cellular adaptation to stress. However, the transcriptional mechanisms linking these pathways remain incompletely understood. Here, we identify Ume6 as a transcriptional repressor that regulates vacuolar protein composition and autophagy in *Saccharomyces cerevisiae* (budding yeast). Dysregulation of Ume6 alters vacuolar protein composition through two distinct mechanisms: increased abundance of multiple vacuolar components and enhanced degradation of selected vacuolar membrane proteins by elevated expression of the vacuolar E3 ubiquitin ligases. *UME6* deletion also increases autophagic activity, whereas *UME6* overexpression delays autophagy induction. Furthermore, we find that Ume6 coordinates with another transcriptional repressor, Rph1, and its regulatory activity is regulated by multiple protein kinases, including Rim15, Rim11, and Mck1. Together, our study reveal a broad transcriptional regulatory mechanism through which Ume6 coordinates vacuolar protein composition and autophagy, providing new insights into the regulation of lysosome-related functions in response to cellular stress.

## Introduction

The lysosome is the catabolic hub of the cell, responsible for maintaining cellular homeostasis through the degradation of macromolecules, recycling of nutrients, and turnover of aged organelles (Ballabio and Bonifacino, 2020; Yang and Wang, 2021). These functions are achieved through the lysosome’s unique composition of lumenal acid hydrolases and wide range of transmembrane transporters and channels. Cellular components, including proteins, nucleic acids, and lipids, are delivered to the lysosome for degradation, and the resulting digestion products are recycled (Yang and Wang, 2021). A major pathway that delivers cellular components to lysosomes is autophagy (“self-eating”), which enables cells to recycle intracellular components and recover nutrients during starvation (Klionsky, 2005). Autophagy begins with the formation of the autophagosome, a double-membrane bound vesicle that originates from the phagophore assembly site (PAS), where cellular cargoes are sorted and engulfed. The phagophore expands to form an autophagosome, which subsequently fuses with the lysosome, delivering its cargo for degradation and recycling (Klionsky, 2007). Consistent with their essential roles in cellular homeostasis, dysregulation of lysosome biogenesis and autophagy has been implicated in numerous human diseases, including cancer and neurodegenerative disorders (Klionsky et al., 2021; Morishita and Mizushima, 2019; Tang et al., 2020; Udayar et al., 2022).

Lysosomal function and autophagy are closely interconnected, particularly during nutrient stress. During starvation, lysosome biogenesis can be stimulated to accommodate the increased demand for intracellular degradation and nutrient recycling (Lakpa et al., 2021). This response accompanies a substantial increase in autophagic flux (Kroemer et al., 2010), and both processes are achieved primarily through transcriptional upregulation of the corresponding genes. Many lysosomal genes, including those encoding lumenal hydrolases and membrane transporters, are transcriptionally regulated under stress conditions (Muller et al., 2015). Likewise, numerous autophagy-related genes (ATGs) are transcriptionally induced in response to environmental and metabolic cues (He and Klionsky, 2009). Conversely, altered expression of lysosome- and autophagy-related genes have been observed in various human disorders (Erie et al., 2015; Fassl et al., 2020; Nixon, 2013), highlighting the importance of transcriptional regulation in maintaining lysosomal function and autophagic activity. Moreover, these two processes must be coordinated because effective autophagy requires not only increased autophagic activity but also an appropriate lysosomal composition to support cargo degradation and nutrient recycling (Yim and Mizushima, 2020). Thus, coordinated regulation of lysosomal composition and autophagy is essential for cellular adaptation to changing environmental conditions.

To date, multiple transcription factors have been identified as regulators of lysosome biogenesis, autophagy, or both. Among them, the best characterized are TFEB and TFE3, which are members of the MiT/TFE family of transcription factors (Steingrimsson et al., 2004). TFEB/TFE3 promote the expression of lysosomal and autophagy-related genes by binding to a conserved DNA sequence known as the “CLEAR” (Coordinated Lysosomal Expression and Regulation) motif (Palmieri et al., 2011). Recently, another DNA-binding protein, upstream stimulatory factor 2 (USF2), has been reported to compete with TFEB to regulate lysosomal and autophagy genes (Kim et al., 2024). These findings highlight the importance of transcriptional networks in coordinating lysosomal function and autophagy. However, in yeast cells, whether there are transcription factors that coordinate vacuole (the yeast equivalent of lysosome) composition and autophagy remain largely unknown.

In this study, we identify Ume6 as a broad transcriptional repressor of both vacuole protein composition and autophagy in yeast cells. Ume6 directly represses genes encoding vacuolar membrane proteins, lumenal hydrolases, protein quality-control machinery, and components of multiple autophagy pathways. Dysregulation of Ume6 leads to multiple downstream effects, including altered vacuolar protein composition and changes in autophagic flux during starvation. Mechanistically, loss of Ume6 alters vacuolar protein composition through two complementary mechanisms: increased expression of target vacuolar proteins and enhanced degradation of other membrane proteins mediated by E3 ubiquitin ligases. Finally, we show that the starvation-associated reduction in Ume6 abundance and promoter occupancy depends on the coordinated functions of Rim15, Rim11, and Mck1. Together, our findings uncover a broad regulatory network linking lysosome composition and autophagy and provide new insight into the transcriptional mechanisms underlying cellular stress adaptation.

## Results

### Atg22 traffics to the vacuole through AP-3 pathway, with both termini facing the cytosol

To investigate, we first focused on Atg22. Atg22 is a vacuolar membrane amino acid exporter which recycles amino acids from autophagic degradation (Sugimoto et al., 2011; Suriapranata et al., 2000; Yang et al., 2006), making it an ideal reporter for studying the coordinated regulation of vacuole biogenesis and autophagy.

To study the regulation of Atg22, we sought out to identify how it is delivered to the vacuolar membrane. In yeast cells, vacuole membrane proteins can be trafficked through two independent pathways: the VPS pathway and the AP-3 pathway (Figure 1A) (Robinson et al., 1988; Stepp et al., 1997). In wild-type (WT) cells, Atg22-GFP localized predominantly to the vacuole membrane (Figure 1B). Deletion of *PEP12* (endosomal t-SNARE) or *VPS27* (ESCRT component) disrupted VPS trafficking, trapping the known VPS cargo Vph1-BFP in small cytosolic vesicles or the class E compartment (Russell et al., 2012), respectively, while Atg22-GFP and the AP-3 cargo Zrc1-mCherry (mCh) remained on the vacuolar membrane (Figure 1B). Deletion of the AP-3 complex component Apl6 caused some Atg22 accumulation in the vacuolar lumen (Figure S1A), implying its degradation when forced into the VPS pathway. Together, these results indicate that Atg22 is trafficked through the AP-3 pathway.

**Figure 1:**
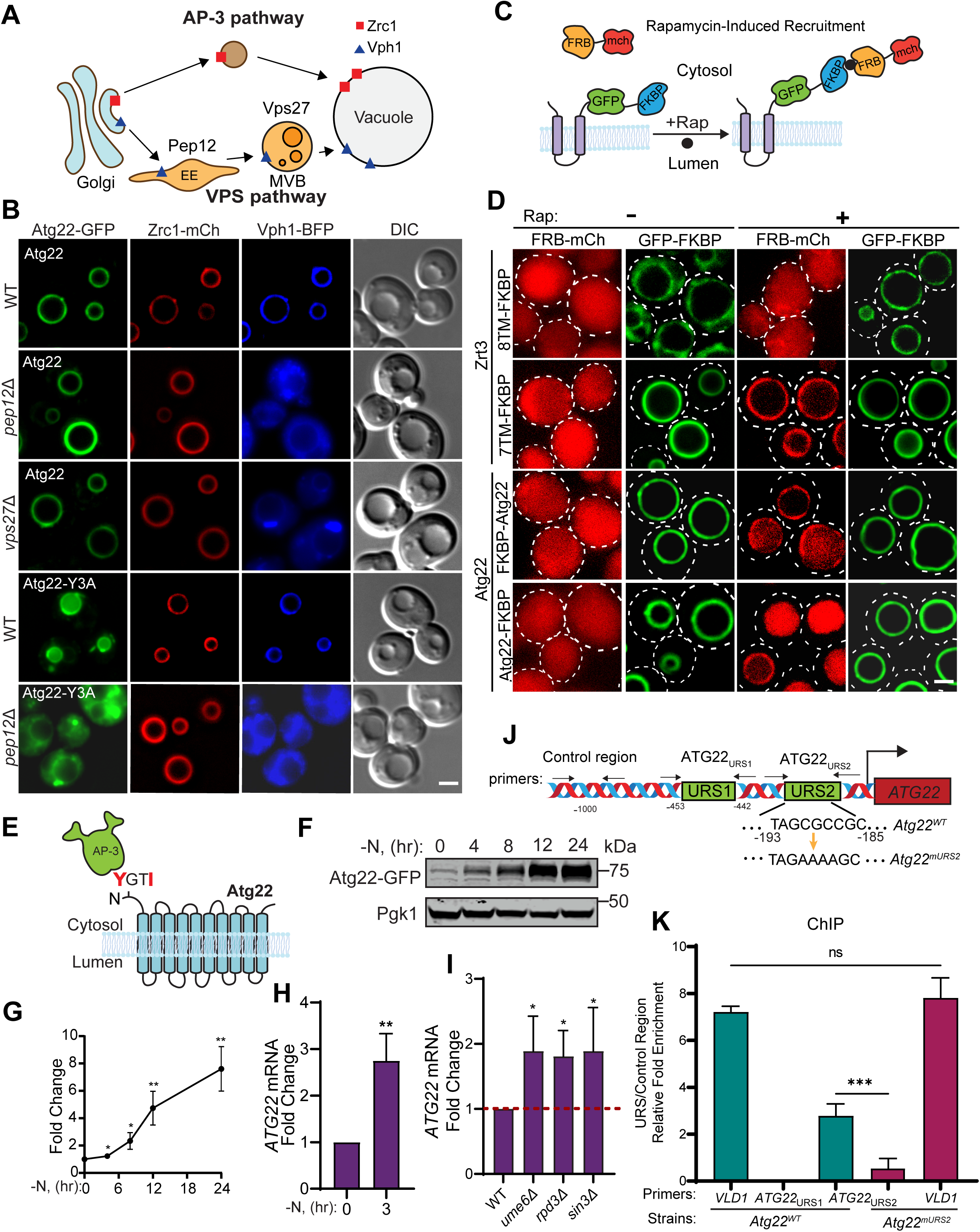
Transcriptional regulation of *ATG22* by the Ume6-Rpd3L complex. **(A)** A cartoon showing the AP3 and VPS pathways for trafficking vacuolar membrane proteins to the vacuole. EE, early endosome; MVB, multivesicular body. **(B)** Subcellular localization of Atg22-GFP (WT and Y3A mutant, both are expressed from a plasmid due to its low endogenous level), Zrc1-mCherry, and Vph1-BFP in WT, *pep12Δ*, and *vps27Δ* strain cells. **(C)** Cartoon depicting the rapamycin-induced recruitment assay. **(D)** FRB-mCherry localization before and after rapamycin treatment in Zrt3^7TM^-GFP-FKBP, Zrt3^8TM^-GFP-FKBP, GFP-FKBP-Atg22, and Atg22-GFP-FKBP strain cells. **(E)** A cartoon showing that AP-3 complex recognizes the tyrosine motif on the cytosolic N-terminus of Atg22. **(F)** Western blot showing the protein level of Atg22-GFP during nitrogen starvation. **(G)** Quantification of (F). **(H)** qPCR showing the mRNA level of *ATG22* after nitrogen starvation. **(I)** qPCR showing the *ATG22* mRNA levels in WT, *ume6*Δ, *rpd3*Δ, and *sin3*Δ strain cells. **(J)** Model showing the design of ChIP assay to test the putative Ume6 binding sites in *ATG22^WT^* and *ATG22^mURS2^*strain cells. **(K)** ChIP-qPCR analysis showing the binding of Ume6 to the *VLD1* and *ATG22* promoter regions in *ATG22^WT^* and *ATG22^mURS2^* strain cells. The error bars represent SD (n = 3 biological repeats). *p ≤ 0.05, **p ≤ 0.01, ***p ≤ 0.001, ****p ≤ 0.00001, student’s t-test. Scale bars, 2 μm.

AP-3 pathway cargoes normally contain either an acidic di-leucine motif ([DE]XXXL[LI]) or a tyrosine-based motif (YXXØ) for their sorting (Braulke and Bonifacino, 2009; Guarnieri et al., 1993). Sequence analysis of Atg22 protein identified two conserved putative tyrosine motifs at residues _3_YGTI_6_ and _46_YIPL_49_ (Figure S1B). Mutating Y3 to alanine caused Atg22^Y3A^-GFP to accumulate in cytosolic puncta and the vacuolar lumen in WT cells (Figure 1B). Further deletion of the *PEP12* gene completely blocked its vacuolar delivery, resulting in the accumulation of cytosolic punctae similar to Vph1-BFP. In contrast, the vacuolar trafficking of Atg22^Y46A^-GFP was not affected (Figure S1A). These results suggest that the motif _3_YGTI_6_ is important for Atg22 trafficking through the AP-3 pathway.

If the _3_YGTI_6_ motif is required for AP-3 sorting, the N-terminus of Atg22 that contains this motif should face the cytosol. However, various topology prediction programs produced conflicting models regarding Atg22 topology (Figure S1C). To resolve this, we first utilized a pHluorin-based assay. pHluorin is a pH-sensitive protein whose fluorescence is quenched in acidic environments such as the vacuolar lumen (Figure S1D). As a control, pHluorin fused to Zrt3, an 8-TM vacuolar transporter with its C-terminus facing the lumen (MacDiarmid et al., 2000), showed no fluorescence. The fluorescence was restored upon vacuolar deacidification by deleting the V-ATPase subunit Vma4 or Bafilomycin A1 treatment, or shifting the pHluorin tag to the cytoplasmic side by deleting the eighth transmembrane helix (Zrt3-7TM, Figure S1D). When tagged with pHluorin at either the N or C terminus, Atg22 showed vacuolar membrane signals, indicating that both termini are exposed to the cytosol.

As a complementary approach, we developed a rapamycin-induced recruitment (RIR) assay (Figure 1C). FRB is fused with mCh and expressed in the cytosol, whereas GFP-FKBP is fused to the protein of interest. Upon addition of rapamycin, FRB and FKBP will undergo dimerization if the tagged terminus faces the cytosol, recruiting the cytosolic FRB-mCh to the GFP-labeled membrane. As proof of principle, cytosol-facing Zrt3-7TM-GFP-FKBP efficiently recruited FRB-mCh, whereas lumen-facing Zrt3-8TM-GFP-FKBP failed to do so (Figure 1D). FRB-mCh was recruited to the vacuole when GFP-FKBP was fused to either terminus of Atg22, confirming that both termini face the cytosol (Figure 1D-E).

### Transcriptional repression of *ATG22* by Ume6-Rpd3 complex

Next, we investigated how Atg22 is regulated under cellular stress. As shown in Figure 1F-G, chromosomally tagged Atg22 protein levels increased dramatically during nitrogen starvation, consistent with previous reports (Suriapranata et al., 2000; Yang et al., 2006). To determine if this upregulation occurs at the transcriptional level, we generated a promoter reporter strain by replacing the coding sequence (CDS) of *ATG22* with GFP (Figure S1F; hereafter referred to as pAtg22-GFP). The protein level of pAtg22-GFP increased during nitrogen starvation (Figure S1G-H), suggesting its transcriptional induction. Consistent with this observation, quantitative real-time PCR (qPCR) analysis showed that *ATG22* mRNA levels increased following nitrogen depletion (Figure 1H).

Bioinformatics analysis revealed two putative upstream repression sequences (URS) for binding of the transcription factor Ume6 on the promoter region of *ATG22* (Figure 1J). Ume6 has been previously shown to coordinate with the Rpd3L histone deacetylase complex to repress gene expression (Kadosh and Struhl, 1997). To investigate the role of Ume6-Rpd3L complex in regulating *ATG22* transcription, we first examined *ATG22* mRNA levels in cells lacking Ume6, Rpd3 (the catalytic unit of the complex), or Sin3 (a major structural component of the complex). As shown in Figure 1I, all three mutants exhibited an elevated level of *ATG22* mRNA compared to WT cells. Consistently, increased pAtg22-GFP levels were observed in these mutants (Figure S1I–J), suggesting that the Ume6–Rpd3L complex functions as a transcriptional repressor of *ATG22*.

Next, to determine whether *ATG22* is a direct target of Ume6, we performed chromatin immunoprecipitation followed by qPCR (ChIP-qPCR). Primer pairs were designed around the two predicted Ume6-binding sites (URS1 and URS2) within the *ATG22* promoter, as well as a control region lacking a putative URS (Figure 1J). *VLD1*, a previously characterized Ume6 target, was included as a positive control (Yang et al., 2018; Yang et al., 2020). Compared with mock samples lacking antibodies (Figure S1K) or a control region lacking a URS (Figure 1K), significant Ume6 enrichment was detected at the predicted URS2 site, but not URS1, in the *ATG22* promoter. Importantly, Ume6 enrichment at the *ATG22* promoter was largely abolished when we mutated the URS2 motif (GCGCC to GAAAA; *ATG22^mURS2^*, Figure 1J-K), while its enrichment at *VLD1* remained unchanged, suggesting that Ume6 binds directly to the promoter of *ATG22* through URS2 motif.

### Ume6 controls vacuolar protein composition through transcription and selective turnover

To determine whether this Ume6-mediated transcriptional repression is a broader mechanism, we performed an RNA-seq in WT and *UME6* deletion cells during mid-log phase growth. Differential gene expression (DGE) analysis revealed significant upregulation of many previously characterized Ume6 target genes (Figure S2A), including *INO1*, *SLZ1*, and *HOP1*, consistent with its established role in repressing meiotic genes (Jackson and Lopes, 1996). Notably, nearly half of the annotated vacuolar genes (41 out of 95) were upregulated in *ume6Δ* cells (Figure S2B), 37 of which contain putative Ume6 binding sequences (Figure 2A, S2B) identified by FIMO (Find Individual Motif Occurrences) analysis (Grant et al., 2011).

**Figure 2:**
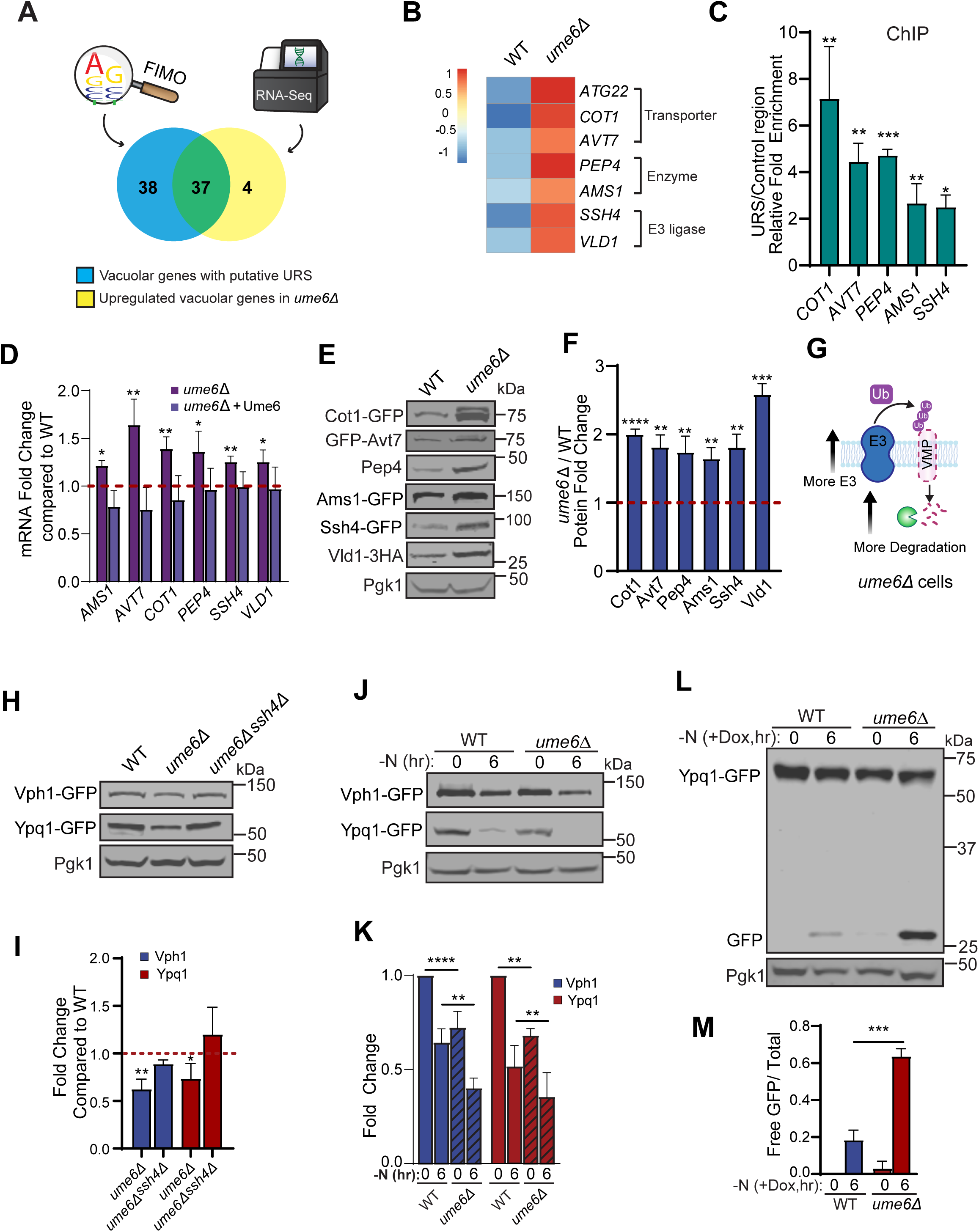
Ume6 negatively regulates the expression of many vacuolar genes. **(A)** Venn diagram showing the overlap of upregulated vacuolar genes after *UME6* deletion with vacuolar genes containing a putative URS for Ume6 binding. **(B)** Heatmap showing the expression change of selected vacuolar genes from RNA-seq data. **(C)** ChIP-qPCR analysis of Ume6 binding to the promoters of selected vacuolar genes. Fold change represents enrichment of binding site region primers over inactive region primers. **(D)** qPCR analysis showing mRNA levels of selected vacuole genes in WT, *ume6Δ*, and *ume6Δ*+*UME6* strain cells. **(E)** Western blot of selected vacuolar proteins in WT and *ume6Δ* strain cells. **(F)** Quantification of (E). **(G)** Cartoon depicting enhanced vacuole membrane protein degradation by E3 ligase upregulation after *UME6* deletion. **(H)** Western blot showing protein levels of Ypq1-GFP and Vph1-GFP in wild-type, *ume6Δ*, and *ume6Δ ssh4Δ* strain cells. **(I)** Quantification of (H). **(J)** Western blots showing the degradation of Ypq1-GFP and Vph1-GFP during nitrogen starvation in WT and *ume6Δ* strain cells. **(K)** Quantification of (J). **(L)** Western blots showing the degradation of TET-off–controlled Ypq1-GFP during nitrogen starvation in WT and *ume6Δ* strain cells. **(M)** Quantification of (L). Dox, Doxycycline. The error bars represent SD (n = 3 biological repeats). *p ≤ 0.05, **p ≤ 0.01, ***p ≤ 0.001, ****p ≤ 0.00001, student’s t-test.

We selected six candidate Ume6 targets for further analysis (Figure 2B), including the vacuole membrane proteins Cot1(MacDiarmid et al., 2000) and Avt7 (Tone et al., 2015), the lumenal hydrolases Pep4 (Ammerer et al., 1986) and Ams1 (Chantret et al., 2003), and the vacuolar E3 ligase complex components Ssh4 (Leon et al., 2008; Li et al., 2015b) and Vld1 (Yang et al., 2018; Yang et al., 2020). For all six genes tested, the enrichment of Ume6 at the predicted URS regions (Figure S2C) was significantly higher than control regions (Figures 2C, S2D), suggesting the direct binding of Ume6. Consistently, the mRNA levels of each target gene were upregulated in *ume6Δ* cells and were restored upon the reintroduction of *UME6* (Figure 2D). Furthermore, *UME6* deletion also resulted in elevated protein levels of each target (Figure 2E-F). Together, these results suggest that Ume6 plays a general role in repressing vacuole biogenesis.

Vacuolar protein composition needs to be precisely regulated through coordinated control of gene transcription and protein degradation. In particular, unneeded membrane proteins are turned over through the ubiquitination of vacuolar E3 ubiquitin ligases (Li et al., 2015a; Li et al., 2015b; Yang et al., 2021; Yang et al., 2020; Zhang et al., 2021). We therefore hypothesized that misregulation of Ume6, which increases the levels of these E3 ligases (Figure 2E), alters vacuolar protein composition by promoting excessive turnover of membrane proteins (Figure 2G). Indeed, the level of two vacuole membrane proteins, Vph1 and Ypq1, both recognized for degradation by Ssh4 (Li et al., 2015a; Yang et al., 2020), were reduced in *ume6Δ* cells (Figure 2H). This reduction was likely caused by elevated Ssh4 levels, as deletion of *SSH4* restored protein abundance. The effects of *UME6* deletion also influenced how vacuole responded to cellular stress. During nitrogen starvation, the levels of both Vph1 and Ypq1 decreased more rapidly in *ume6Δ* cells (Figure 2J-K), accompanied by increased free GFP accumulation (Figure S2E–G; uncropped images corresponding to Figure 2J), indicating enhanced degradation. Lastly, to confirm the accelerated degradation is from the preexisting protein pool, we expressed Ypq1-GFP under the control of a TET-OFF system, in which *YPQ1* transcription is suppressed by adding doxycycline. As shown in Figure 2L–M, the degradation of Ypq1-GFP occurred substantially faster in *ume6Δ* cells.

Together, these results suggest that Ume6 is a general regulator of vacuole composition. Loss of Ume6 alters vacuolar composition through two complementary mechanisms: direct upregulation of Ume6 target genes and indirect downregulation of other vacuolar proteins caused by elevated E3 ligases.

### Ume6 represses *ATG* genes and negatively regulates multiple autophagy pathways

Our RNA-seq analysis also identified that many autophagy-related genes (*ATG*s; 20 out of 31 examined), including the known Ume6 targets *ATG8* (Bartholomew et al., 2012) and *ATG32* (Aihara et al., 2014), were upregulated in *ume6Δ* cells (Figures 3A, S3A), with more than half containing putative URS motifs. To investigate if Ume6 broadly represses autophagy genes, we selected six candidates from distinct autophagy pathways: *ATG8* (core autophagy machinery) (Xie et al., 2008), *ATG19* and *ATG34* (cytosol to vacuole targeting pathway, Cvt) (Scott et al., 2001; Suzuki et al., 2010), *ATG32* and *ATG33* (mitophagy) (Kanki and Klionsky, 2010; Okamoto et al., 2009), and *ATG40* (ER-phagy) (Mochida et al., 2015) (Figure S3B). It is worth noting that DGE analysis showed only a modest 1.2-fold upregulation of *ATG40* in *ume6Δ* cells, making its change less apparent on the heatmap (Figure S3A). Nevertheless, we included *ATG40* as a candidate because of its important role in ER-phagy.

**Figure 3:**
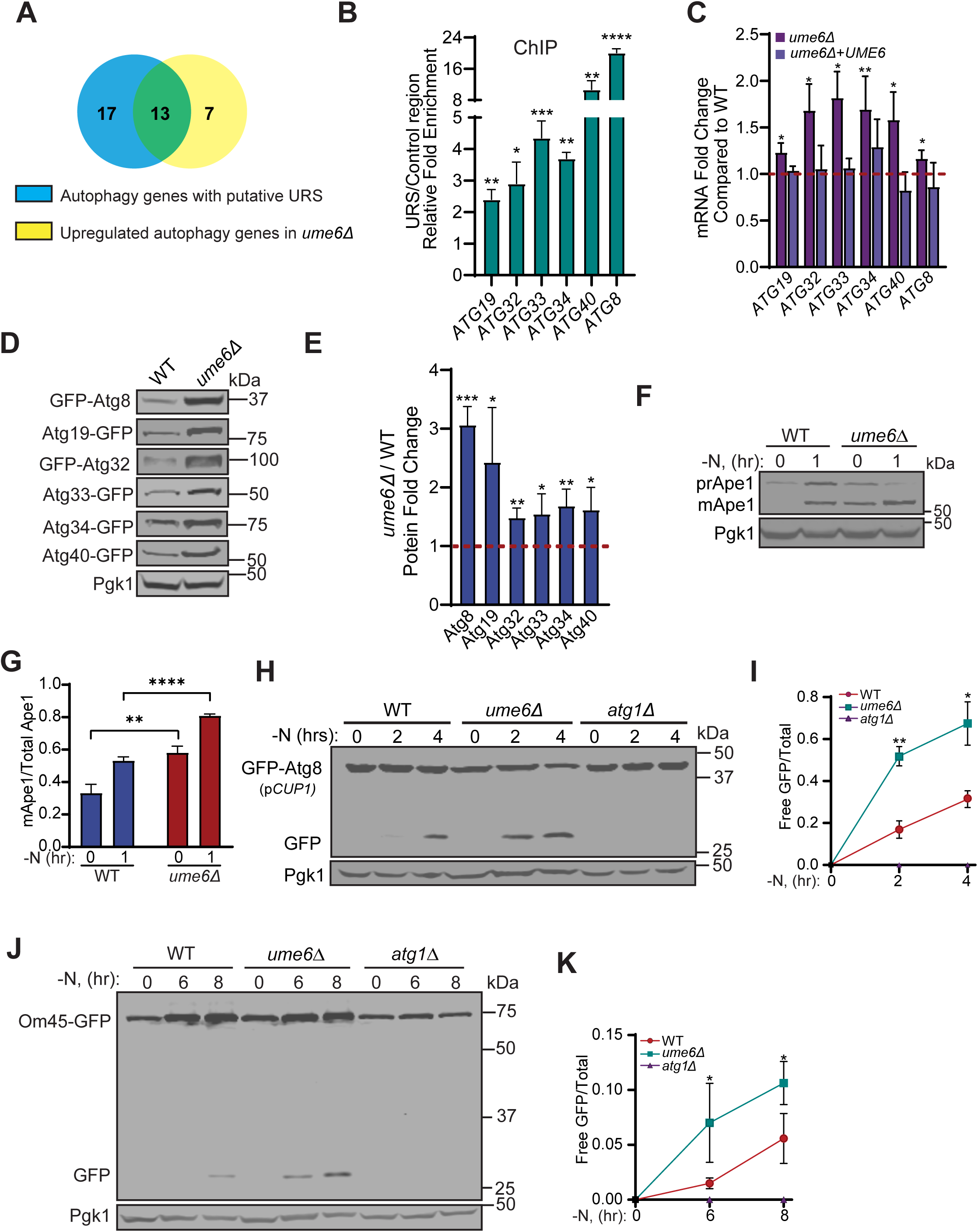
Ume6 represses several *ATG* genes and negatively regulates autophagy. **(A)** Venn diagram showing the overlap of upregulated autophagy (*ATG*) genes after *UME6* deletion with *ATG* genes containing a putative URS for Ume6 binding. **(B)** ChIP-qPCR analysis of Ume6 binding to the promoters of selected autophagy genes. Fold change represents enrichment of the binding site region primers over inactive region primers. **(C)** qPCR analysis showing mRNA levels of selected *ATG* genes in WT, *ume6Δ*, and *ume6Δ*+*UME6* strain cells. **(D)** Western blot of autophagy proteins in WT and *ume6Δ* strain cells. **(E)** Quantification of (D). **(F)** Western blot showing processing of prApe1 into mature Ape1 during nitrogen starvation in WT and *ume6Δ* cells. **(G)** Quantification of (F). **(H)** Western blot showing levels of GFP-Atg8 in WT, *ume6Δ*, and *atg1Δ* strain cells during nitrogen starvation. **(I)** Quantification of (H). **(J)** Western blot showing the levels of Om45-GFP in WT, *ume6Δ*, and *atg1Δ* strain cells during nitrogen starvation. **(K)** Quantification of (H). The error bars represent SD (n = 3 biological repeats). *p ≤ 0.05, **p ≤ 0.01, ***p ≤ 0.001, ****p ≤ 0.00001, student’s t-test.

ChIP-qPCR confirmed the enrichment of Ume6 at the predicted URS-containing regions of all six promoters (Figures 3B, S3C), indicating the direct binding of Ume6. Furthermore, the mRNA levels of each gene were upregulated in *ume6Δ* cells and restored upon reintroduction of *UME6* (Figure 3C). Finally, we examined the protein levels of GFP-tagged target genes in both WT and *ume6Δ* cells and found that the protein levels of all targets were increased in *ume6Δ* cells (Figure 3D–E).

To determine whether Ume6 can regulate autophagy activity, we first examined the Cvt pathway by monitoring Ape1 processing. Upon Cvt-mediated delivery to the vacuole, precursor Ape1 (prApe1) is proteolytically processed into mature Ape1 (mApe1) (Stromhaug and Klionsky, 2001; Suzuki et al., 2002) (Figure 3F–G). *ume6Δ* cells exhibited increased Ape1 processing under both nutrient-rich and starvation conditions, suggesting enhanced Cvt pathway activity. We next monitored GFP-Atg8 processing for general autophagic flux (Klionsky et al., 2007). Because endogenous *ATG8* is a target of Ume6 and is upregulated after *UME6* deletion, we expressed GFP-Atg8 from a plasmid driven by the *CUP1* promoter. Under the *CUP1* promoter, the expression level of GFP-Atg8 was lower in *ume6Δ* cells, but still showed a higher processing rate during starvation, suggesting increased autophagic flux in *ume6Δ* cells (Figure 3H-I). Similarly, higher mitophagy activity was observed in *ume6Δ* cells when we monitored the degradation of Om45-GFP (Figure 3J-K), a mitochondrial outer membrane protein that is commonly used as a mitophagy reporter (Kanki et al., 2009). Together, all these results suggest that Ume6 negatively regulates autophagy activity.

### Ume6 cooperates with distinct transcriptional repressors to control autophagy

Previous studies have identified several repressors of autophagy pathway, including Pho23 (Jin et al., 2014), and Rph1 (Bernard et al., 2015). Our identification of Ume6 as another broad repressor of *ATG* genes prompted us to investigate their relationships. Interestingly, Pho23 is also a component of the Rpd3L complex (Loewith et al., 2001). As shown in Figure 4A-B, the *ume6Δ pho23Δ* double mutant did not display an additive effect compared with either single mutant, consistent with Ume6 and Pho23 functioning in the same complex. It is noteworthy that Atg9 is a reported target of Pho23 and Rpd3, but not of Ume6 (Figure S3A) (Jin et al., 2014). Consistent with this, *ume6Δ* cells did not exhibit increased Atg9 protein levels, suggesting that the Rpd3–Pho23 complex may regulate *ATG9* independently of Ume6, likely through other DNA-binding factors. In contrast, Atg8 levels were higher in *ume6Δ* cells than in *pho23Δ* cells or double mutant, although the mechanism remains unclear.

**Figure 4:**
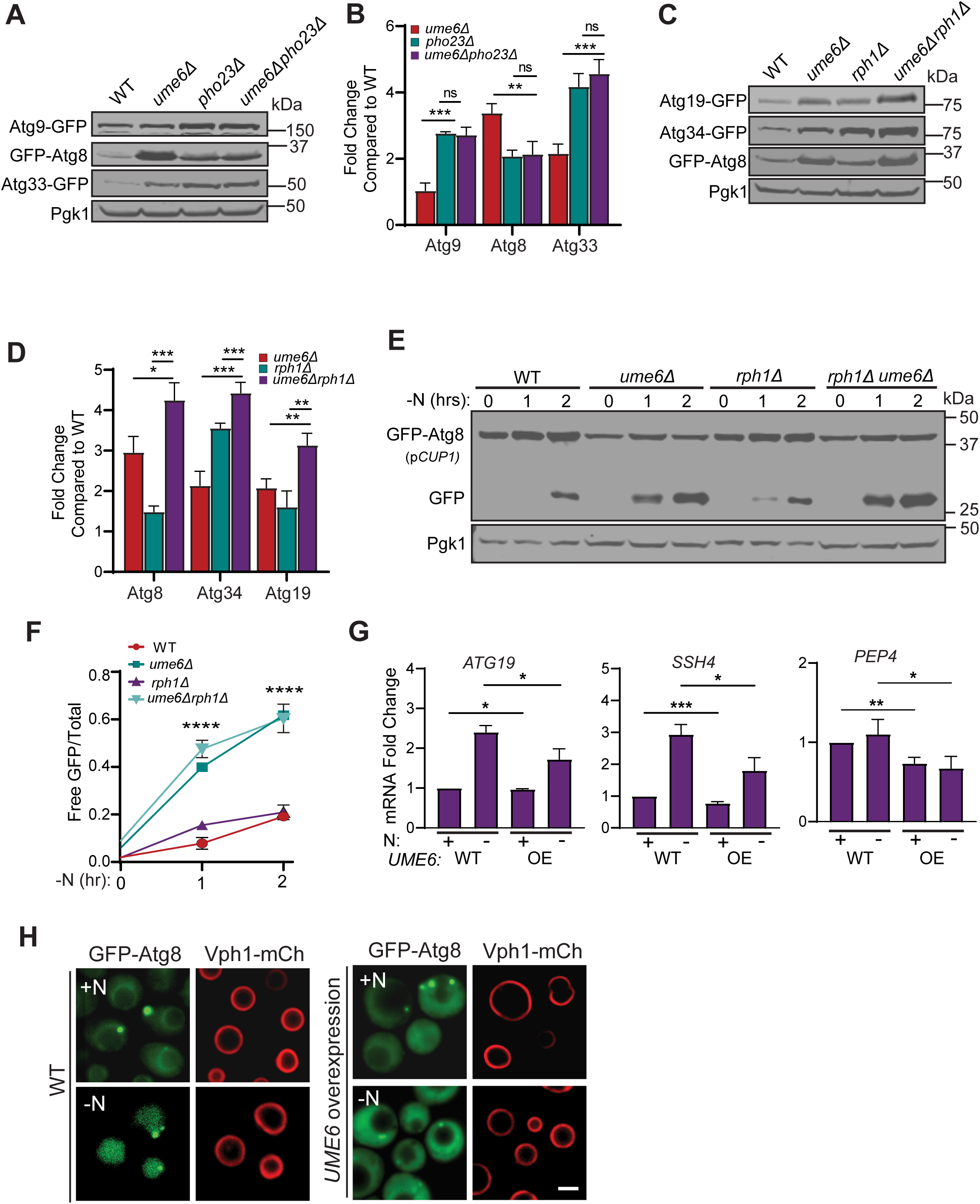
Genetic interactions of Ume6 and effects of Ume6 overexpression. **(A)** Western blot showing the protein levels of Atg9-GFP, GFP-Atg8, and Atg33-GFP in WT, *ume6Δ*, *pho23Δ,* and *ume6Δ pho23Δ* strain cells. **(B)** Quantification of (A). **(C)** Western blot showing the protein levels of Atg19-GFP, GFP-Atg8, and Atg34-GFP in WT, *ume6Δ*, *rph1Δ,* and *ume6Δ rph1Δ* strain cells. **(D)** Quantification of (C). **(E)** Western blot showing the GFP-Atg8 level in WT, *ume6Δ*, *rph1Δ,* and *ume6Δ rph1Δ* strain cells during nitrogen starvation. **(F)** Quantification of (E). Statistical significance was determined using a one-way ANOVA **(G)** qPCR analysis of *ATG19*, *SSH4*, and *PEP4* under nitrogen starvation in WT and *UME6* overexpression cells. **(H)** Subcellular localization of endogenous GFP-Atg8 in WT and *UME6* overexpression strain cells before and after nitrogen starvation. Error bars represent SD (n = 3 biological repeats). *p ≤ 0.05, **p ≤ 0.01, ***p ≤ 0.001, ****p ≤ 0.00001, student’s t-test. Scale bars, 2 µm.

Rph1 is a DNA-binding protein that represses autophagy gene expression through binding to the consensus STRE (stress response element) motif (Nordberg et al., 2014). Our analysis identified multiple *ATG* gene promoters that contain both URS and STRE motifs (Figure S3A), suggesting that Ume6 and Rph1 may coordinately regulate *ATG* gene expression. As shown in Figure 4C-D, Atg8, Atg19, and Atg34 were all upregulated in both *ume6Δ* and *rph1Δ* single mutant cells. Moreover, the double mutants exhibited even higher levels of all three proteins. To determine whether this additive effect also influences autophagy activity, we measured GFP-Atg8 processing (Figure 4E-F). Compared to WT cells, both *ume6Δ* and *rph1Δ* single mutant showed increased GFP-Atg8 processing after 1 hour of starvation, suggesting more rapid autophagy induction. After 2 hours of starvation, *ume6Δ* cells still displayed elevated autophagy activity, whereas the enhanced activity relative to WT in *rph1Δ* cells became attenuated. Consistently, the *ume6Δ rph1Δ* double mutant showed exhibited higher GFP-Atg8 processing than either single mutant during early starvation, but its phenotype mimicked *ume6Δ* during prolonged starvation. Together, these results suggest that Rph1 coordinates with Ume6 in regulating autophagy activity.

### Overexpression of Ume6 inhibits target gene expression and autophagy induction

To further test the role of Ume6 in vacuole biogenesis and autophagy, we analyzed the expression of several Ume6 target genes in WT and *UME6* overexpression cells. As shown in Figure 4G, overexpression of *UME6* caused a modest reduction of all three representative target genes under growth conditions, including *SSH4* (vacuole membrane), *PEP4* (vacuole lumen), and *ATG19* (autophagy), and this repression became more pronounced after 1 hour of starvation. We next investigated whether *UME6* overexpression can alter autophagy activity by examining the localization of endogenously tagged GFP-Atg8. In WT cells, GFP-ATG8 mainly diffused in the cytosol as well as localized to cytosolic punctae in growth conditions (Figure 4H). After 1 hour of nitrogen starvation, GFP-Atg8 was internalized into the lumen of Vph1-mCh-labeled vacuole, indicating the induction of autophagy. In contrast, in cells overexpressing *UME6*, GFP-Atg8 largely remained in the cytosol, suggesting that autophagy induction was delayed.

### Multiple kinases coordinate in regulating Ume6 and downstream gene expression

Lastly, we attempted to identify the upstream signaling pathways for Ume6 regulation. To investigate, we endogenously tagged Ume6 with 3V5 in WT cells and monitored its protein levels during starvation. As shown in Figure 5A-B, Ume6-3V5 levels decreased upon starvation. Consistently, ChIP-qPCR revealed significantly reduced Ume6 enrichment at target genes after starvation (Figure 5C), likely to reflect the Ume6 degradation associated with transcriptional release. Supporting this hypothesis, the starvation-induced decrease of Ume6 levels was largely blocked in the proteasome mutant *cim3-1*(Figure 5D-E).

**Figure 5:**
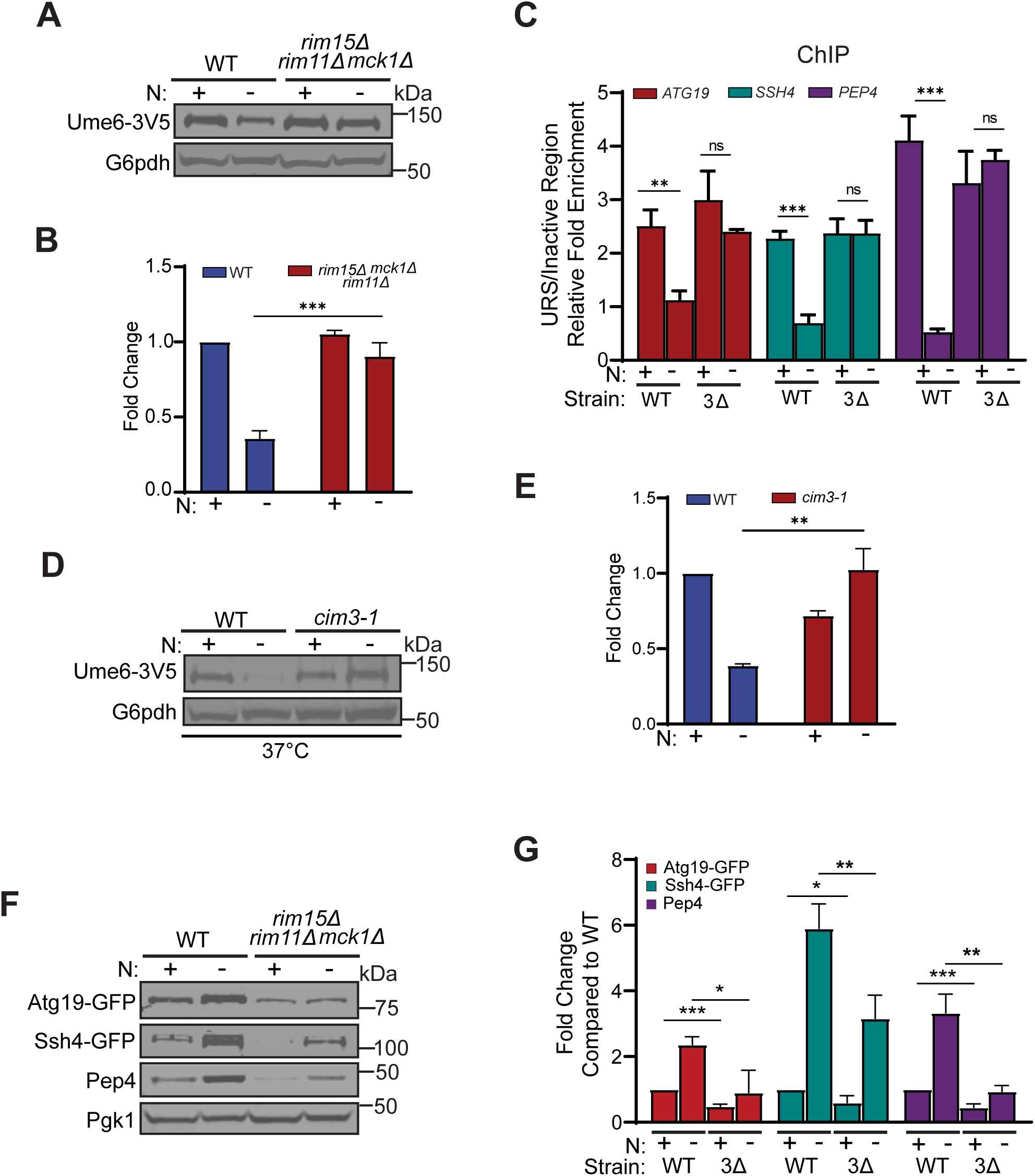
Coordinated regulation of Ume6 and target genes by Rim15, Mck1, and Rim11. **(A)** Western blot showing levels of Ume6-3V5 before and after nitrogen starvation in WT and *rim15Δ rim11Δ mck1Δ* strain cells**. (B)** Quantification of (A). **(C)** ChIP-qPCR analysis of *ATG19*, *SSH4*, and *PEP4* during nitrogen starvation in WT and *rim15Δ rim11Δ mck1Δ* strain cells. Fold change represents enrichment of the binding site region primers over inactive region primers. **(D)** Western blot of levels of Ume6-3V5 during nitrogen starvation in WT and *cim3-1* cells. **(E)** Quantification of (D). **(F)** Western blot showing the protein levels of Atg19-GFP, Ssh4-GFP, and Pep4 in WT and *rim15Δ rim11Δ mck1Δ* strain cells during nitrogen starvation. **(G)** Quantification of (F). The error bars represent SD (n = 3 biological repeats). *p ≤ 0.05, **p ≤ 0.01, ***p ≤ 0.001, ****p ≤ 0.00001, student’s t-test.

Three kinases have previously been implicated in Ume6 phosphorylation: Rim15 (Bartholomew et al., 2012), Rim11, and Mck1 (Aihara et al., 2014; Xiao and Mitchell, 2000). To determine whether these kinases contribute to Ume6-mediated regulation, we monitored Ume6 protein levels in WT, *rim15Δ*, and *mck1Δ rim11Δ* double mutant cells. We chose to use *mck1Δ rim11Δ* double knockout as it has been reported that Rim11 and Mck1 have overlapping functions (Xiao and Mitchell, 2000). As shown in Figure S4A-B, *rim15Δ* cells exhibited unchanged basal Ume6-3V5 levels but substantially delayed Ume6 degradation during starvation. In contrast, *mck1Δ rim11Δ* cells showed elevated basal Ume6-3V5 levels under nutrient-rich conditions, while Ume6 levels still decreased significantly upon starvation. Notably, in *rim15Δ rim11Δ mck1Δ* triple mutant, both the starvation-induced decrease in Ume6 protein levels and its dissociation from target gene promoters were largely blocked (Figure 5A-C), highlighting the roles of these three kinases in regulating Ume6.

Consistent with these observations, when we examined the expression of Ume6 target genes in the same mutant cells, target protein levels were comparable to WT in *rim15Δ* cells under growth conditions, whereas their starvation-induced upregulation was delayed (Figure S4C-D). In *mck1Δ rim11Δ* cells, the basal levels of all three target proteins examined were reduced, likely due to elevated Ume6 levels. Moreover, their induction during starvation was largely unaffected, with the exception of Atg19. In *rim15Δ rim11Δ mck1Δ* triple mutant cells, both basal expression and starvation-induced upregulation of target proteins were strongly repressed (Figure 5F-G). Together, these results suggest that multiple kinases, including Rim15, Rim11, and Mck1, coordinately regulate Ume6 and downstream gene expression. Among them, Rim11 and Mck1 appear to be more responsive in regulating Ume6 under nutrient-rich conditions, whereas Rim15 plays a crucial role in stress response.

## Discussion

### Ume6 is a general regulator coordinating autophagy and vacuolar remodeling

In this study, we identify Ume6 as a general repressor that coordinates vacuolar protein composition and autophagy in yeast cells (Figure 6). Under nutrient-rich conditions, Ume6 directly binds to the promoters of its target genes and represses their expression. During starvation, Ume6 dissociates from these promoters, relieving transcriptional repression and enabling the upregulation of genes involved in vacuolar function and autophagy. On the autophagy side, multiple *ATG* genes involved in distinct autophagy pathways, including non-selective autophagy, the Cvt pathway, ER-phagy, and mitophagy, are upregulated to support diverse forms of autophagy. On the vacuole side, remodeling involves both increased expression of selected proteins, such as the lumenal hydrolase Pep4 and vacuolar membrane protein Cot1, and the selective turnover of others, such as the vacuolar membrane protein Ypq1. The degradation process is promoted by the upregulation of Ume6-controlled E3 ligases, including Ssh4. Through this complementary regulation of protein synthesis and degradation, Ume6 regulates vacuolar composition to meet changing cellular demands, including those associated with increased autophagic activity.

**Figure 6:**
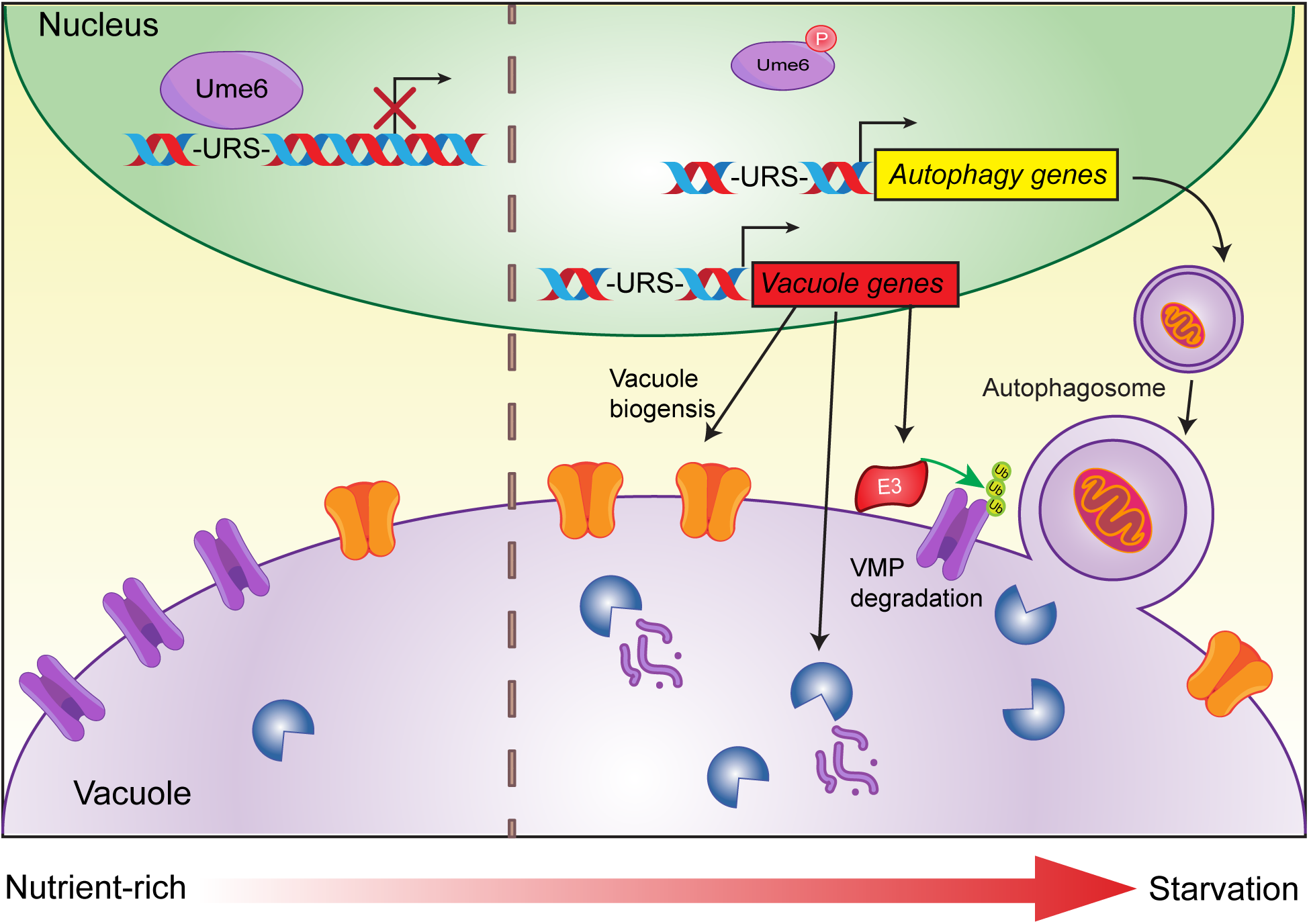
A model summarizing the transcriptional repression of lysosome remodeling and autophagy by Ume6. For details, please see the Discussion section.

As two major cellular stress-response pathways, autophagy and its destination organelle, the lysosome, must be coordinately controlled. In mammalian cells, TFEB and TFE3 function as master regulators that control the expression of hundreds of genes involved in the autophagy– lysosomal pathway, metabolism, and immune responses (Sardiello et al., 2009). More recently, USF2, has been shown to compete with TFEB for repressing autophagy and lysosomal genes (Kim et al., 2024). In yeast cells, several general transcription factors have been identified in autophagy regulation, including Pho23 and Rph1 as transcriptional repressors, and Gcn4 and Gln3 as transcriptional activators (Bernard et al., 2015; Hinnebusch and Natarajan, 2002; Jin et al., 2014; Minehart and Magasanik, 1991). However, the transcriptional regulation of vacuolar protein composition remains less well understood, and how vacuolar remodeling is coordinated with autophagy has not been fully elucidated. Our identification of Ume6 as a broad transcriptional repressor of vacuolar and autophagy-related genes provides a framework for understanding how these processes are coordinated at the transcriptional level.

Ume6, named for Unscheduled Meiotic gene Expression, is required for the repression of early meiotic genes such as *INO1* (Jackson and Lopes, 1996; Strich et al., 1994). In budding yeast, meiosis is initiated by nutrient depletion, upon which Ume6-mediated repression is relieved through replacement by the transcriptional activator Ime1 (Kassir et al., 1988). Regulation by Ume6 is critical for meiotic progression, as deletion of *UME6* leads to significant defects in the expression of early meiosis-specific genes (Steber and Esposito, 1995). Here, we identify an additional role for Ume6 in coordinating vacuolar protein composition and autophagy. Given the complexity of the yeast vacuole and autophagy pathways, which involve more than one hundred proteins, it remains to be determined whether dedicated transcriptional activators also coordinate the expression of genes involved in these two pathways and, if so, how they interact with Ume6 to regulate their coordinated expression.

### The coordination between Ume6 and other transcription factors in autophagy regulation

Rph1 has been reported to repress the transcription of several autophagy genes, including *ATG7*, *ATG8*, *ATG9*, *ATG14*, and *ATG29*. Bernard et al. demonstrated that deletion of *RPH1* increases the basal expression levels of these genes, whereas overexpression of Rph1 has the opposite effect. The identification of Ume6 as another broad transcriptional repressor of autophagy genes, together with the observation that these two repressors share many autophagy-related targets, prompted us to investigate the relationship between Ume6 and Rph1. Our results indicate that these two repressors have additive effects on target gene repression. The autophagy genes we tested were upregulated in both single mutants and showed even stronger upregulation in the double mutant. These findings suggest that Ume6 and Rph1 make distinct contributions to the repression of overlapping target genes, potentially through independent or partially independent mechanisms. More interestingly, our data suggest that Rim15 plays an important role in regulating Ume6 and its target gene expression. Notably, the same kinase, Rim15, has also been shown to regulate Rph1 activity (Bernard et al., 2015). This raises the possibility that Rim15 coordinates multiple downstream transcriptional repressors to shape the autophagic response to nutrient stress.

Why do cells need two different repressors on the same promoters, particularly when both are regulated by the same upstream signaling pathway? One possibility is that multiple repressors allow cells to fine-tune transcriptional responses according to cellular needs. For example, we observed that although *rph1Δ* cells display higher autophagy activity during the early stages of starvation, this difference is rapidly diminished as WT cells catch up during prolonged starvation, consistent with the previously described “jump-start” effect (Bernard et al., 2015). In contrast, Ume6 appears to play a more sustained and substantial role in autophagy regulation. These observations suggest that Ume6 and Rph1 may contribute differently to the temporal control of autophagy, with Rph1 having a more prominent effect during early starvation and Ume6 contributing to sustained repression. Such functional specialization could enable cells to modulate the timing and magnitude of autophagy according to the duration and severity of nutrient stress.

It is also notable that these two repressors seem to have different target preferences, as their deletions have distinct effects on different genes. For example, Atg19 and Atg8 are more regulated in *UME6* deletion cells, whereas deletion of *RPH1* has a greater impact on *ATG34* expression. In addition, the reported binding motif for Rph1, 5’-TTAGGGG-3’, overlaps with the consensus stress response element (STRE), which is also recognized by the transcription factor Msn2 (Jacquet et al., 2003). Msn2 functions as a general stress-response factor involved in the regulation of heat shock, oxidative stress, and starvation responses (Kuang et al., 2017; Sadeh et al., 2011; Yamamoto et al., 2008). There is also evidence linking Msn2 to autophagy regulation through control of the core autophagy factor Atg8 (Vlahakis et al., 2017). However, whether Msn2 plays a broader role in the regulation of autophagy has not yet been explored, and whether Msn2 also functionally interacts with Ume6 and Rph1 also remains an open question. Together, these findings suggest that the transcriptional regulation of autophagy is highly complex, and that we are only beginning to understand the underlying regulatory network.

## Materials and Methods

### Yeast strains, plasmids, media, and growth conditions

All yeast strains used in this study are listed in Table S1. Strains were constructed using the lithium acetate-based transformation protocol (Gietz and Woods, 2006). All plasmids used are listed in Table S2 (Longtine et al., 1998). All yeast strains were grown at 26°C, unless indicated otherwise, in either YPD or YNB media before further analysis.

### Starvation assays

Yeast cells were grown to mid-log phase (OD_600_∼0.5–0.7). For nitrogen starvation, cells were pelleted at 3,500 rpm for 5 minutes. Cells were then washed with nitrogen starvation medium (YNB without amino acids and ammonium sulfate, with 2% glucose), then incubated in nitrogen starvation medium at 26°C for the remainder of the assay. For rapamycin induced starvation, 0.5 µM rapamycin was added to cultures and incubated at 26°C for the remainder of the assay. For TET-OFF assays, 2 mg/ mL doxycycline was added at a 1:1000 dilution and shaken at 26°C for thirty minutes before being transferred to nitrogen starvation media with doxycycline for the remainder of the assay.

### Chromatin Immunoprecipitation (ChIP)

Chromatin immunoprecipitation was performed based on (Aparicio et al., 2004), with some modifications. Yeast cultures were grown to mid-log phase (OD_600_∼0.5–0.7). For nitrogen starvation experiments cells were first spun down, washed once with YNB-nitrogen media, and resuspended in YNB-nitrogen media for the indicated time. Formaldehyde to 1% was added under stirring for crosslinking DNA/proteins. Glycine (125 mM) was used to quench crosslinking, and cells were lysed with bead vortexing for 45 minutes 4°C. Chromatin was sheared via sonication to 250-500 bp fragments and immunoprecipitated with protein G magnetic beads (Fisher, 10003D) and anti-V5 antibody (1:300, Fisher R960-25), excluding the mock sample which included beads but no antibody, overnight. DNA-protein complexes were eluted and crosslinks reversed for 6 hours with proteinase K digestion at 65°C. DNA was purified and extracted via phenol-chloroform purification and ethanol extraction. Eluted DNA was analyzed via qPCR wherein primers were designed for the active region (promoter containing URS motif) and an inactive region (upstream of promoter not containing URS motif). IP Ct for active and inactive regions were normalized to input Ct, and relative fold enrichment was calculated based on the fold change difference between the active and inactive regions. For mock Ct calculations, IP Ct was subtracted from Mock Ct, and the fold change was calculated. Primers are listed in Table S3.

### RNA isolation and qRT-PCR

Yeast cells were grown in YPD to mid-log phase (OD_600_∼0.5–0.7) and 14 OD_600_ unit of cells were collected. Total RNA was extracted from pellets using TRIzol (145105; Life Technologies) and PureLink RNA Mini Kit (1938678; Invitrogen). Approximately 1 µg of RNA was used for cDNA synthesis using Verso cDNA Synthesis Kit (AB1453A; Fisher) with anchored oligo (dT) primers. qPCR was done using the Power SYBR Green PCR Master Mix (1708558D; Thermo Fisher Scientific) with the primers targeting either *UBC6* (housekeeping gene) or specific genes listed in Table S3. For each sample, the relative transcript levels were determined by normalizing them to *UBC6* levels.

### RNA-Seq

Total RNA samples were prepared via RNA-Isolation method. cDNA libraries were constructed and sequenced at Novogene in Davis, Oregon. Sequencing was performed on an Illumina NovaSeq X Plus system, producing stranded paired-end reads of 150 bp to a minimum depth of 30 million paired-end reads. The raw sequences were aligned to the *S. cerevisiae* reference genome obtained from Ensemble and quantified using Salmon (v 1.10.2). For each sample, the identified transcripts were quantified without normalization (raw counts). Differential expression analysis was performed with Deseq2 (v 1.50.2). Putative binding sites were analyzed for gene promoter regions using FIMO (v 5.5.8). The raw data of RNA-seq is available on NCBI through accession # PRJNA1466526.

### Microscopy and image processing

Microscopy and image processing were done with Leica Thunder Wide Field Fluorescence system (Leica Microsystems). Yeast cells were grown to mid-log phase (OD_600_∼0.5–0.7) and washed with MilliQ water, then imaged at room temperature (RT). For the bafilomycin and pHluorin studies, cells were incubated for 1 hour with 20 nM bafilomycin A1 at 26°C before being washed with MilliQ water and imaged at RT. For rapamycin recruitment assays, cells were incubated with 10 µM rapamycin for 1 hour at 26°C before being washed with MilliQ water and imaged at RT. Image acquisition and deconvolution were performed with the LAX program. The images were further cropped and adjusted by using ImageJ (National Institutes of Health).

### Sample preparation for Western blotting and antibodies

Yeast cells were treated with 10% TCA and kept at 4°C for at least 12 hours. After washing with 0.1% TCA, cell pellets were dissolved in 2× boiling buffer (50 mM Tris pH 7.5, 1 mM EDTA, and 1% SDS) and 100 µL of glass beads were added. Samples were vortexed at RT for 5 minutes to lyse cells and heated at 65°C for 5 minutes. Equivalent amount of 2× urea sample buffer (150 mM Tris, pH 6.8, 6 M urea, 6% SDS, 40% glycerol, 100 mM DTT, and 0.01% bromophenol blue) was added and samples were vortexed and then heated at 65°C for 5 minutes each. The supernatants were collected, and approximately 1 OD_600_ unit of cells were loaded in each lane for SDS-PAGE and transferred to nitrocellulose membranes for blotting. The following antibodies were used in this study: Pgk1 (1:10,000, 22C5D8; Invitrogen), rabbit anti-GFP (1:3000, TP401; Amsbio), mouse anti-3HA (1:3000, 901503; BioLegend), and mouse anti-V5 (1:3000, Fisher; R960-25). Antibody against Pep4 and Ape1 was a generous gift from Dr. Daniel J. Klionsky (University of Michigan).

### Western blotting quantification

The Western blot images were scanned using the Licor Odyssey DLx system, and band intensities were quantified from the raw data files using the Image Studio Ver5.2 software.

### Statistical Analysis

All experiments were independently performed at least three times. The values are presented as the mean ± SD, n= biological repeats. The significance of differences between groups was evaluated using unpaired two-tailed Student’s t tests for comparisons involving two groups. For multiple group comparisons one way ANOVA was used. The P value of <0.05 was considered statistically significant.

## Data availability statement

The raw data of RNA-seq is available on NCBI through accession # PRJNA1466526. Authors declare that data supporting the findings of this study are available within the article and its supplementary information files or from the corresponding author upon request.

## Acknowledgements

We thank members of the Yang laboratory for their technical support. We are grateful to Dr. Daniel J. Klionsky for sharing reagents. This research is supported by a startup fund from the University of Denver, and a NIH grant R35GM159962 to X. Yang.

## Author contributions

Conceptualization, O.Evans., and X.Yang.; Methodology, O.Evans., A. Hoy., I. Woo., and X.Yang.; Investigation, O.Evans., A. Hoy., I. Woo., and X.Yang.; Original Draft, O.Evans., and X.Yang.; Editing, X.Yang.; Funding Acquisition & Supervision, X.Yang.

